# Chitosan films for microfluidic studies of single bacteria and perspectives for antibiotics susceptibility testing

**DOI:** 10.1101/591602

**Authors:** Julie Tréguier, Loic Bugnicourt, Guillaume Gay, Mamoudou Diallo, Salim Islam, Alexandre Toro, Laurent David, Olivier Théodoly, Guillaume Sudre, Tâm Mignot

## Abstract

Single cell microfluidics is powerful to study bacteria and determine their susceptibility to antibiotics treatment. Glass treatment by adhesive molecules is a potential solution to immobilize bacterial cells and perform microscopy but traditional cationic polymers such as poly-lysine deeply affect bacterial physiology. In this work, we chemically characterized a class of chitosan polymers for their biocompatibility when adsorbed to glass. Chitosan chains of known length and composition allowed growth of *Escherichia coli* cells without any deleterious effects on cell physiology. Combined with a machine-learning approach, this method could measure the antibiotics susceptibility of a diversity of clinical strains in less than 1 hour and with higher accuracy than current methods. Last, chitosan polymers also supported growth of *Klebsiella pneumoniae*, another bacterial pathogen of clinical significance. The low cost of chitosan slides and their simple implementation makes them highly versatile for research as well as clinical use.

## Introduction

In recent years, microfluidics coupled with live-cell imaging have revolutionized bacteriology, testing directly the impact of rapid and controlled environmental transitions on cell physiology. With the advent of super-resolution microscopy, the bacterial cell can now be further explored at unprecedented resolution, tracking cellular processes one molecule at a time *(1)*. The impact of these methods is not limited to basic research because single cell approaches are unquestionably powerful at determining antimicrobial susceptibility (AST, Antimicrobial Susceptibility Testing) in record time *(2, 3)*.

However, a major technical bottleneck with the implementation of single cell approaches for super-resolution or AST, is the immobilization of bacterial cells. Agar surfaces have been widely used and support the growth of a wide range of bacterial species. However, this method has several limits:

i. Agar surfaces are not compatible with high-end microscopy (HEM) methods that require cells to adhere to glass, for example Total Internal Reflection Fluorescence Microscopy (TIRFM) and all single molecule microscopy techniques (PALM, STORM, STED).
ii. Because the adhesion of bacterial cells to Agar surfaces is generally weak, these surfaces cannot be manipulated in aqueous environments and the experimental conditions are generally set by diffusion through the agar substrate. However, this approach does not allow rapid changes of the medium or injection of chemicals and thus the kinetics and precise dose-dependent effects are poorly controlled *(3, 4)*.

Alternative methods have remedied these issues by growing bacteria immediately in contact with a glass surface. Because most bacteria do not directly adhere to glass, immobilization procedures are required, which include direct physical immobilization of the bacteria in micro channels or glass functionalization by adhesive polymers. The use of micro-channels is certainly compatible with HEM and it allows fast AST with high accuracy *(2, 5)*. However, this method requires expert handling, complex nanolithography to produce the channels and extensive development to be used for the study of a given bacterial species. Alternatively, bacterial adhesion on glass can be obtained by functionalizing a glass slide with adhesive polymers/molecules. This approach can also be difficult because the polymer must be fully biocompatible and the functionalization procedure and surface chemistry can be complex. Indeed, although this approach has been widely used for eukaryotic cells, the choice for polymers biocompatible with bacteria is limited. Cationic polymers such as poly-lysine bind glass surfaces effectively and promote adhesion of a wide range of bacterial species. However, poly-lysine also generates cell envelope stress and has been shown to dissipate/diminish the membrane potential in several Gram negative of Gram positive species (e.g. *Escherichia coli* or *Bacillus subtilis (6–8)*). For clinical microbiology applications, this issue is particularly sensitive because changes in the membrane potential can directly affect antimicrobial susceptibility *(9)* and thus produce false negative or even worst, false positive results in AST. Thus, there is a need in developing new functional polymers with neutral effects on bacterial physiology for single cell AST studies.

This work originated from the observation that chitosan polymers can support bacterial adhesion and motility on surfaces (in the case of *Myxococcus xanthus* and *Bacillus subtilis (1)*). Herein, we investigated if chitosan-treated glass slides could also support bacterial growth when inserted in commercial microfluidic chambers. Using a specific chitosan polymer (with a high degree of acetylation) and a new controlled functionalization procedure, we showed that Chitosan-Coated Slides (CCS) can support the growth of *E. coli* during multiple generations without any effects on bacterial fitness. Using clinical *E. coli* strains obtained from intestinal and urinary tract infections ((I/U)TI) of known antibiotics susceptibilities, we showed that CCS allowed fast direct determination of AST. Last, CCS can be derived to promote growth of other so-called ESKAPE pathogens such as *Klebsellia pneumoniae*, which also raise significant problems for antibiotics treatment *(10)*. We conclude that chitosan-based functionalization procedures are promising for their application in bacterial single cell studies for basic research but also potentially, in clinical contexts.

## Results

### Functionalizing glass slides with chitosan polymers

Chitosan is a linear polysaccharide composed of randomly distributed β-(1→4)-linked D-glucosamine and N-acetyl-D-glucosamine units (Figure S1A). Its physicochemical properties are highly dependent on its macromolecular parameters (i.e. average molar mass *M_w_*, and degree of acetylation DA). A control of these parameters is needed to ensure robustness when studying the physicochemical and biological behavior of chitosan polymers. Indeed, growth and motility where not always reproducible when glass slides were coated with raw commercial chitosan and this variability could be due to the poor chemical characterization of commercial stocks, which contain chains of variable DA, molar mass and statistical distribution of the acetyl groups.

Consistent with this, size exclusion chromatography (SEC-MALLS/RI) analysis performed on chitosan from a commercial source (Sigma-Aldrich, See Methods), revealed an important dispersity in polymer chains length (Polydispersity Index Đ = 2.65). It is essential to control the dispersity of chitosan chains because slight variations in molar mass and DA of chitosan polymers can be associated to a wide range of biological responses: cell adhesion, wound healing and even bacterial stasis and lysis *(11, 12)*. As a general trend, it is mandatory to determine and control each molecular parameter in order to understand their impact on bacterial physiology and to ensure reproducibility of our experiments.

To this aim, we first generated a large library of chitosan polymers with various DA and molar masses *(13)* (Chito-library). Different molar masses (*M_w_*, 180 kg/mol and high *M_w_*, 460 kg/mol) were obtained by selecting chitosan from different source (shrimp or squid). To control the acetylation levels, the polymer chains were re-acetylated *in vitro* to produce DAs of 1%, 10%, 15%, 25%, 35%, 45%, and 55% *(14)*. Each polymer was characterized by SEC to control its molar mass and by ^1^H NMR to measure its DA (Figure S1A, see Methods).

Flat and homogeneous layers of polymer in the nanometer range (i.e. 15 nm < thickness < 150 nm) were obtained by Spin-coating of chitosan solutions with controlled concentration and pH (Figure 1A). Substrates were chosen to be either silicon wafers for physicochemical characterization, or boro-silicate glass coverslips for bacterial single cell studies (Figure S1B, no incidence of the substrate was noticed on the physicochemical characteristics of chitosan layers). The thickness, uniformity, wettability and morphology of chitosan ultrathin films prepared from the chito-library were systematically examined by ellipsometry, tensiometry, optical microscopy, profilometry or atomic force microscopy (Figure 1B). Whatever the formulations studied, thickness and wettability of chitosan layers were highly reproducible (e.g. 23.3 ± 1.3 nm, and 37.8 ± 1.2°, for chitosan formulation of DA 55%, [c] = 0.67%, *M_w_* = 180 kg/mol, with n = 10), with a Root Mean Square (RMS) roughness lower than 1 nm. The detailed information on the physicochemical properties of the chitosan thin films will be described in a specialized dedicated publication.

**Figure 1.**
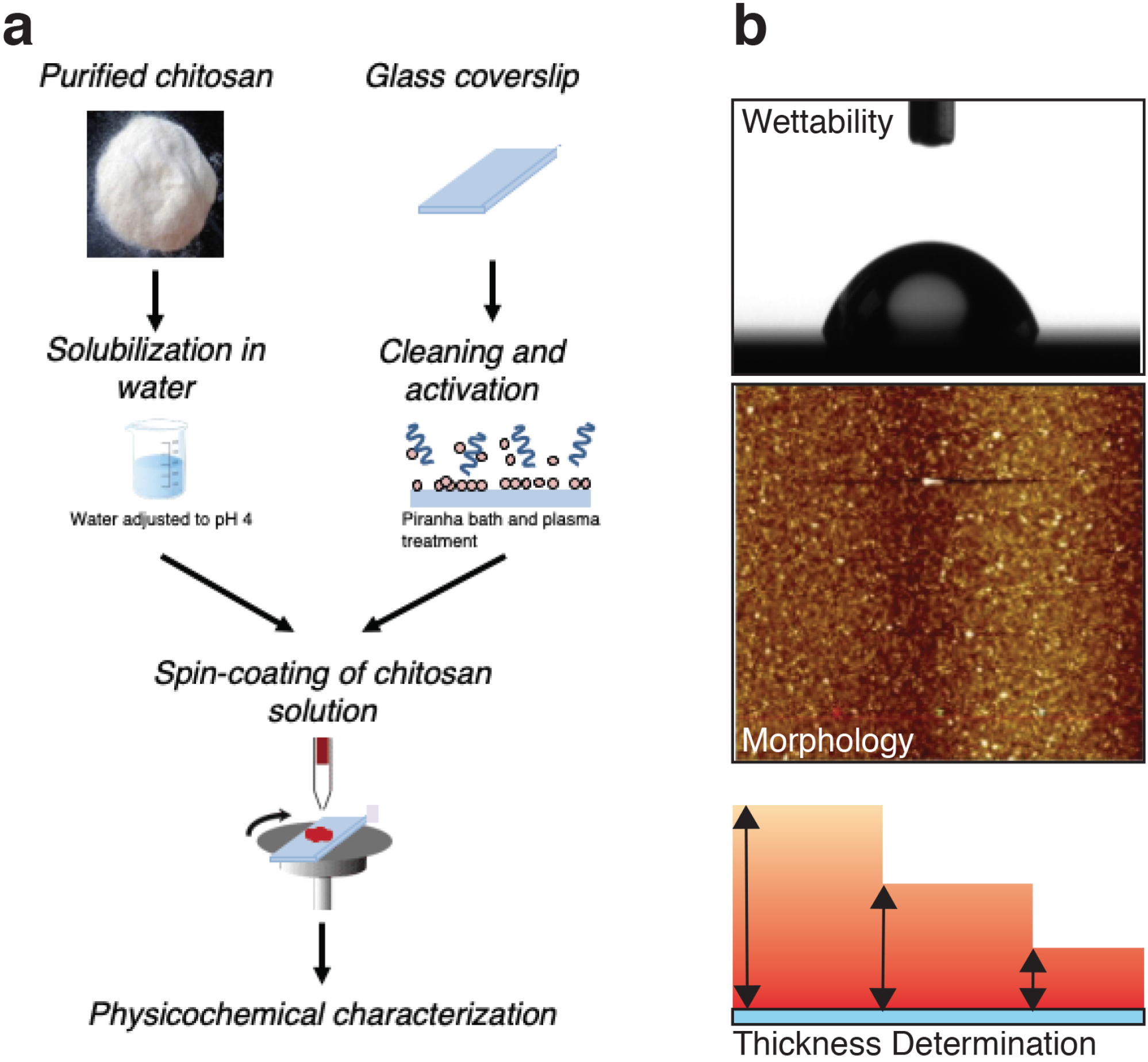
Functionalization of glass slides with Chitosan polymers. **(a)** Protocol for glass surface modification and characterization of chitosan layers (See Methods for details). **(b)** Chemical characterization of Chitosan thin films. Thin films are characterized by wetting measurements, morphology imaging (AFM) and their thickness was determined by ellipsommetry (See Methods for details).

We thus successfully generated homogeneous CCS of known polymer molar mass, DA and thickness. By varying the chitosan macromolecular parameters and chitosan solution characteristics, more than 50 different chitosan coatings were thus prepared to be screened for their ability so support bacterial proliferation.

### Specific chitosan polymers promote adhesion and normal growth of *E. coli* cells

We next tested the ability of the various types of CCS to support the adhesion and ultimately growth of the main laboratory *E. coli* K12 strain. To perform this screening, we divided our CCS library in nine representative subclasses, based on source, DA and additional treatments (Table 1). Each CCS type was then mounted at the bottom of a microfluidic cassette and tested for *E. coli* adhesion and growth (see Methods).

We found that while LB-grown *E. coli* cells did not adhere to uncoated glass slides, they adhered to all CCS types, showing that chitosan can indeed promote adhesion of *E. coli*. However, while *E. coli* cells did generally proliferate on these surfaces, growth was frequently abnormal, evidenced by cell filamentation and morphological aberrations (Figure S2A). Nevertheless, one type of CCS obtained with chitosan polymers of DA 55%, M_W_ of 156 kDa and thickness of 32 nm supported normal growth (C5, Figure 2A, Table 1). To further characterize this chitosan class, we tested 156 kDa polymers of varying DAs and found that DAs ≥50% were required for biocompatibility (Table 1). In addition, formulation was important because acid rinsing negatively impacted the biocompatibility of the procedure (Table 1).

**Figure 2.**
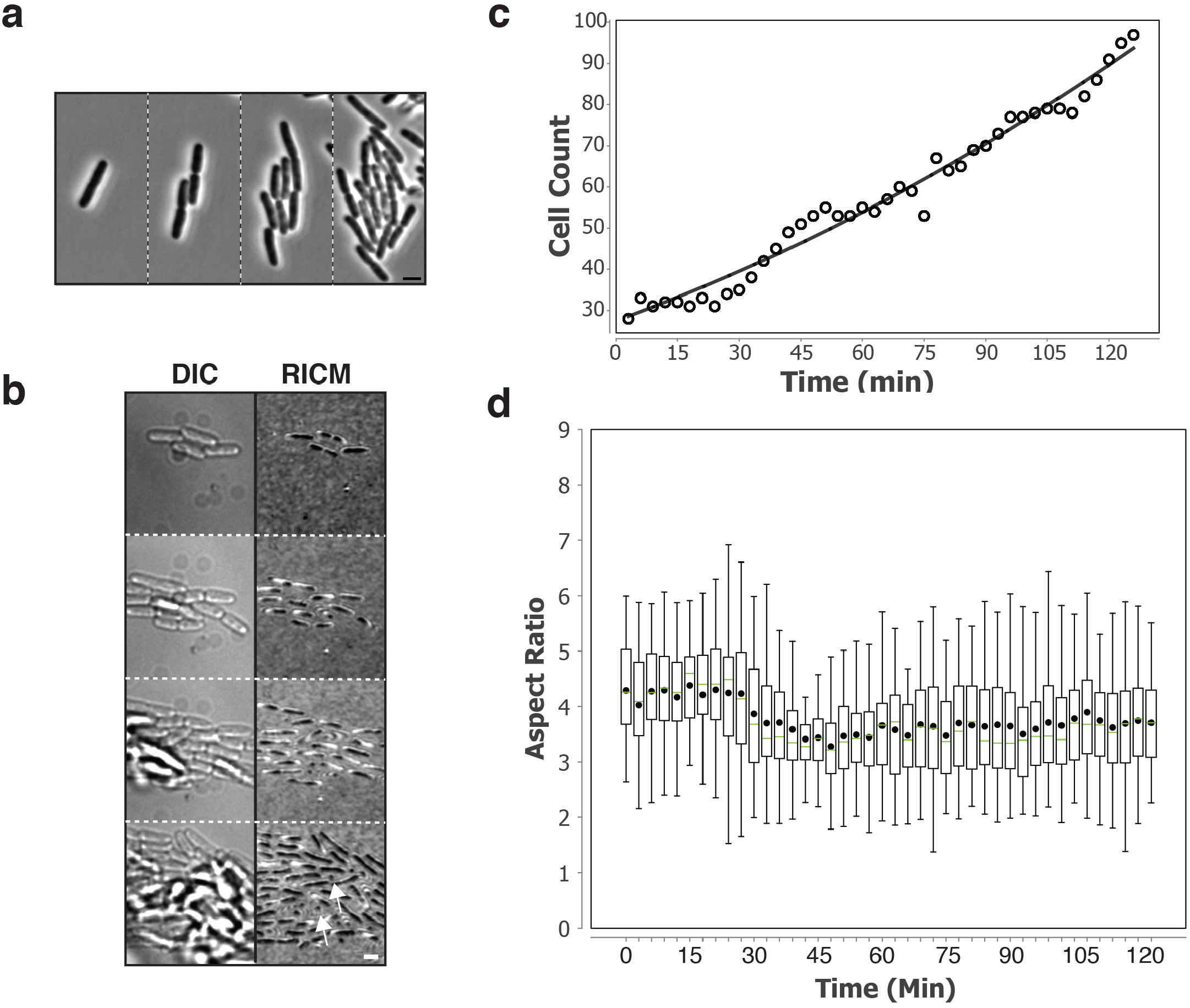
Growth of E. coli on selected C5 Chitosan slides. **(a)** Growth of *E. coli* K12 on C5. Shown are snapshots separated by 54 min after growth initiation (left panel). See associated Supplemental movie S1 for the full time-lapse. Scale bar = 2 μm. **(b)** Adhesion of *E. coli* on C5 as measured by Reflection Interference Contrast Microscopy (RICM). *E. coli* is shown by Nomarski (DIC, left) revealing the three-dimensional organization of the *E. coli* micro-colony and by RICM to reveal the adhesion sites (observed as dark areas, right). Note that the cells remain tightly adhered to the Chitosan surface even at the latest time points when the micro-colony clearly expands above the focal plane. White arrows point to areas where the cells remain adhered by the cell pole only, allowing them to grow away from the Chitosan surface. See associated Supplemental movie S2 for the full time-lapse. Scale bar = 2 μm. **(c)** Growth of *E. coli* K12 on C5. Shown is an exponential fit of the number of cells as a function of time. **(d)** Morphology of *E. coli* on C5 over time. The aspect ratio are determined from phase contrast images of adhered cells and correspond to the ratio between the lengths of the long axis and the short axis of the cell.

We next characterized the ability of C5 to promote adhesion and growth in detail. *E. coli* K12 formed mono-layered micro-colonies and could be monitored for up to 6 generations after which the cells started growing above the focal plane defined by the glass slide. Expansion of the *E. coli* micro-colony in 3D could occur because tight adhesion of the monolayer forced the daughter cells to grow away from the immediate surface, which has been shown to act as driving force for bacterial colony and biofilm development *(15)*. To test this possibility, we analyzed *E. coli* cell adhesion to C5 by Reflection Interference Contrast Microscopy (RICM), a technique that allows imaging of intimate cell contacts with glass surfaces *(16)*. RICM revealed that each cell remained in close contact with the glass surface by adhering along their axis. Surface escape was due to steric constraints and vertical growth of bacteria adhered via their cell poles (Figure 2B, Movie S2). In contrast, when we performed RICM on a CCS type that created abnormal cell torsions, it was apparent that the dividing cells only adhered via the cell poles, explaining cell detachment and the emergence of torsions (Figure S2B). Consistent with the RICM results, *E. coli* cells remained attached to C5 event when subjected to shear stress of up to 12 dyn/cm^2^ (which is comparable to shear stress generated in aorta *(17)*, see Methods).

We next tested whether C5 created detectable stress on K12 *E. coli* growth. *E. coli* K12 cells grew exponentially with a generation time similar to the generation of *E. coli* grown under agitation in liquid culture at room temperature (Figure 2C, C5 experiments were conducted at 25°C. C5 also supported growth of *E. coli* at 37°C but all described experiments were performed at 25°C to avoid the use of a thermo-controller system). Cell morphology, measured by the aspect ratio (length/width) remained stable over time, showing that it was not affected on C5 (Figure 2D). Last, to test whether C5 generates long term cellular defects, we allowed *E. coli* cells to develop on C5 until they reached stationary phase and became quiescent for 3 days. These cells resumed growth normally after fresh medium was injected, showing that long term exposure to C5 does not affect cell viability (Figure S2C). We conclude that C5 is a well-adapted chitosan to grow *E. coli* K12 cells on glass surface in microfluidics chambers.

### CCS allow fast Antibiotics Susceptibility Testing (AST)

Beyond their obvious use in research applications, CCS could provide a fast and reliable tool for AST. For this, CCS should be significantly faster and at least as reliable as currently used methods. To test this, we incubated *E. coli* K12 on C5 and injected Ampicillin, which rapidly resulted in the typical cell elongation and formation of a bulge in the septal zone that precludes cell lysis (Figure 3A, Movie S3). The approximate time-to-death was ~120 min (Td), consistent with the kinetics described in other single cell experiments *(18)*. On C5, Ampicillin generates the same cellular defects as in other studies and could thus be used for AST.

**Figure 3.**
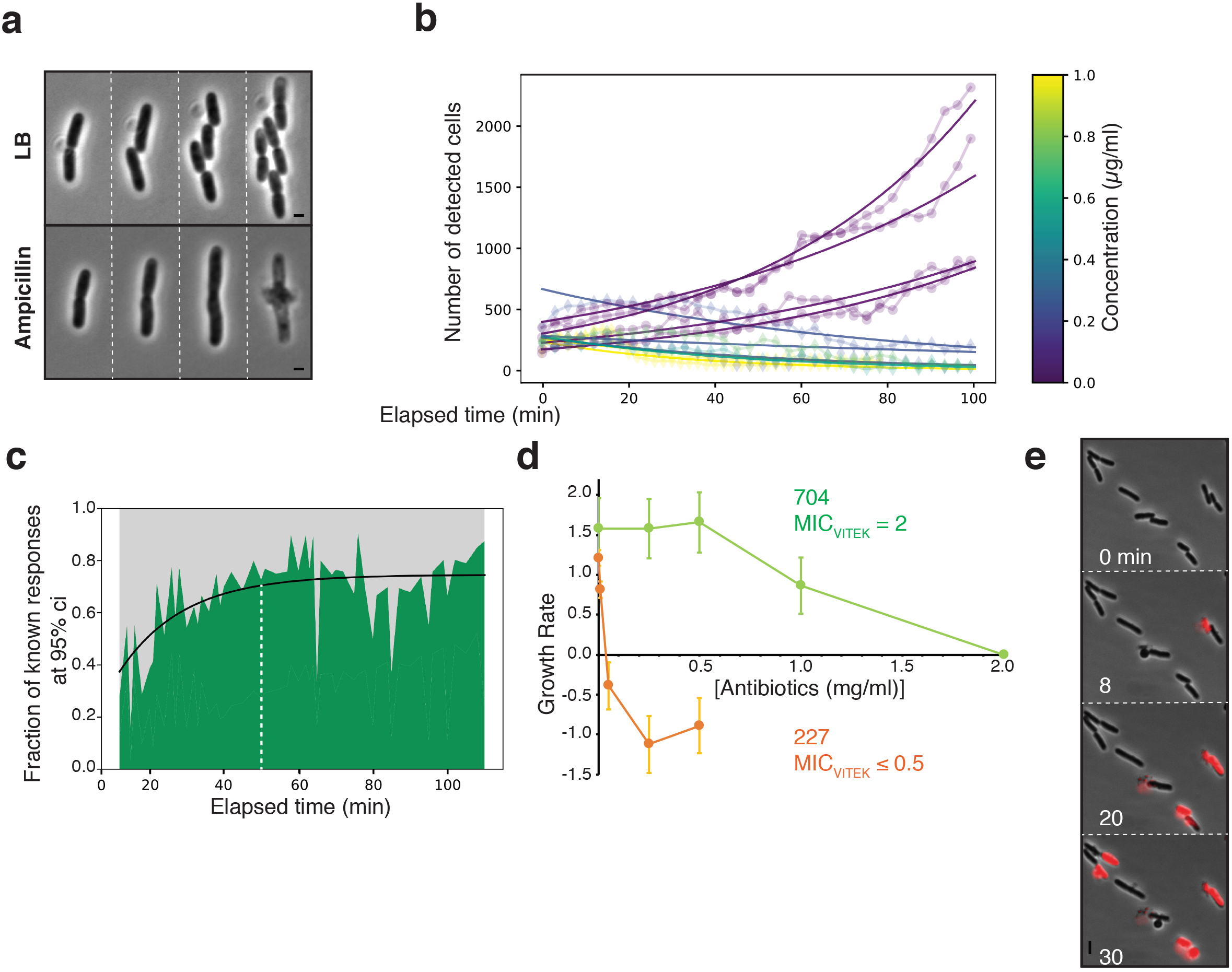
C5-CCS can be used for fast AST of E. coli clinical strains. **(a)** Ampicillin treatment is effective on C5. Note the characteristic Ampicillin-induced morphological transitions, cell elongation and the formation of a septal lytic “bubble” that precludes cell death. See associated supplemental movie S3 for the full-time lapse. **(b)** Trained detection of Ertapenem effects on growth of *E. coli* clinical strains. Measured growth curves for strain UTI227 with varying concentrations of ETP. Fitted growth curves computed from the number of detected cells across time are color-coded with respect to ETP concentration. For each curve, the plot symbol is circular if the cells survive, and diamond shaped if the cell population stalls or shrinks due to cell death. **(c)** Estimation of the minimal diagnostic time. We performed an estimation of the growth rate for varying time spans for all assays and determined for each time span the fraction of assays for which the response could be ascertained with a 95% confidence interval. **(d)** Comparison of the MICs as measured on CCS with MICs obtained at the hospital. The MIC is determined for growth rates ≤ 0 obtained at given antibiotics concentrations. Note that the hospital (Vitek) and CCS-determined MICs for Mecillinam are similar for UTI704, but that the CCS method measures MICs as low 0.05 for UTI227 in the presence of Ertapenem. **(e)** Detection of cell death by Propidium-Iodide (PI) staining. PI only stains the bacterial DNA of permeable dead cells, which fast and sensitive quantification of MBCs. See associated supplemental movie S4 for a typical time lapse.

We next tested whether antibiotic susceptibility may be determined in less time than the measured Td (as detected by irreversible cell lysis). Indeed, although *E. coli* cells lyse after 2hrs, the action of ampicillin is first characterized by abnormal cell elongation (Figure 3A). Thus, early detection of abnormal cell morphologies would provide a fast method to assess the action of Ampicillin. To do this reliably and computationally, we designed a machine-learning based morphometric method that discriminates abnormal cell morphologies from WT cell morphologies and detects the effect of antibiotics at different treatment times (Figure S3A-C and Methods). Briefly, following segmentation and determination of cell contours, this method allows the direct counting of cells with normal morphologies and thus the determination of growth curves. This approach could readily determine growth curves of an *E. coli* strain isolated from a urinary tract infection and treated with increasing doses of Ertapenem, a relatively large-spectrum Carbapenem standardly used at the hospital (UTI227, Figure 3B). Lethal Ertapenem effects could be detected as early as 50 minutes after addition of the antibiotic with 95% confidence (Figure 3B-3C). Thus, combined with our computational detection method, CCS, and here specifically C5, is a promising tool for fast AST.

### CCS can be used to measure the Minimal Inhibitory Concentration of clinical *E. coli* isolates

To test the potential clinical application of CCS more broadly, we next obtained a collection of 15 clinical isolates derived from Urinary Tract (UTIs, 14 isolates) and Intestinal Tract Infections (ITI,1 isolate) and tested their ability to grow on C5. We determined that 70% of the clinical strains adhered and grew normally on C5 but that this number could be improved to 85%, if the thickness of C5 were increased to 66 nm (Table1), showing that thickness is another important parameter to increase the application spectrum C5 to most *E. coli* clinical strains.

In current clinical practice, the antibiotic susceptibility of a given bacterial strain is determined by its so-called Mimimum Inhibitory Concentrations (MIC), which corresponds to the lowest antibiotic concentration that prevents growth. To test if MICs determined on C5 can be directly compared to MICs determined by standard methods, we further selected two clinical strains of known MICs (as determined by Vitek2, Biomérieux) for Mecillinam (UTI704 MIC=2) and Ertapenem (UTI227 MIC≤0.5) and measured their MICs on C5, extracting growth rates with our computational methods (Figure 3D). In both cases, the results showed remarkable consistency with the Vitek method and in fact, the CCS method was more sensitive allowing to determine that the UTI227 Ertapenem MIC is between 0.01 and 0.05 mg/ml (Figure 3D, Table 2). To further test the validity of the method, we tested the consistency of the measurements over various range of Ertapenem concentrations for UTI227 and showed that its Mecillinam MIC on C5 also matches the Vitek-determined MIC (Table 2). Further MIC measurements on additional clinical strains UTI687 and UTI698 on Ofloxacine and Mecillinam, respectively also showed good consistency with Vitek measures (Table 2). In conclusion, CCS appears a promising tool to measure MICs rapidly and accurately in hospitals.

### CCS can be used to measure the Minimal Bactericidal Concentration of antibiotics

The MIC does not measure microbial death *per se* and thus it cannot distinguish bactericidal from bacteriostatic effects. This can be problematic for treatment, especially since it was discovered that antibiotic treatment can induce bacterial persistence, a seemingly dormant state that could be associated with chronic infections *(19)*. Using the Minimum Bactericidal Concentration (MBC), the lowest antibiotic concentration resulting in bacterial death, would in general be more appropriate but it is a highly time-consuming procedure because it requires regrowing the bacteria after antibiotic treatment. However, as we show above, the CCS technology allows direct observation of *E. coli* cell lysis in the presence of Ampicillin and is therefore a potential tool to determine the MBC of an antibiotic directly and rapidly (Figure 3A). In addition, the microfluidic environment of CCS allows detection of cell death, for example using dyes such as Propidium Iodide (PI) that only bind the bacterial DNA if the bacterial membrane is irreversibly altered. The use of PI can improve detection sensitivity, especially if the detection method is automated. Indeed, addition of PI on *E. coli* treated-Ertapenem allowed cell death detection on C5, suggesting that this method could be used to determine MBCs in clinical contexts (Figure 3E, Movie S4).

### CCS can be adapted to promote surface growth of *Klebsiella pneumoniae*

Next, we were interested in testing whether C5 could be useful to study other clinically-relevant pathogens. In particular, and along with UTI *E. coli* strains, *K. pneumoniae* is a member of the so-called ESKAPE pathogens, characterized by the high resistance of clinical strains to antimicrobial compounds and thus a growing concern in hospital environments *(10)*. Indeed, *K. pneumoniae* could readily grow on C5 (but at 66 nm thickness), with normal morphology and generation time (~40 min, Figure 4A-B, Movie S5). Thus, C5 is a versatile substratum for bacterial adhesion, and could be used in hospitals for AST of ESKAPE pathogens.

**Figure 4.**
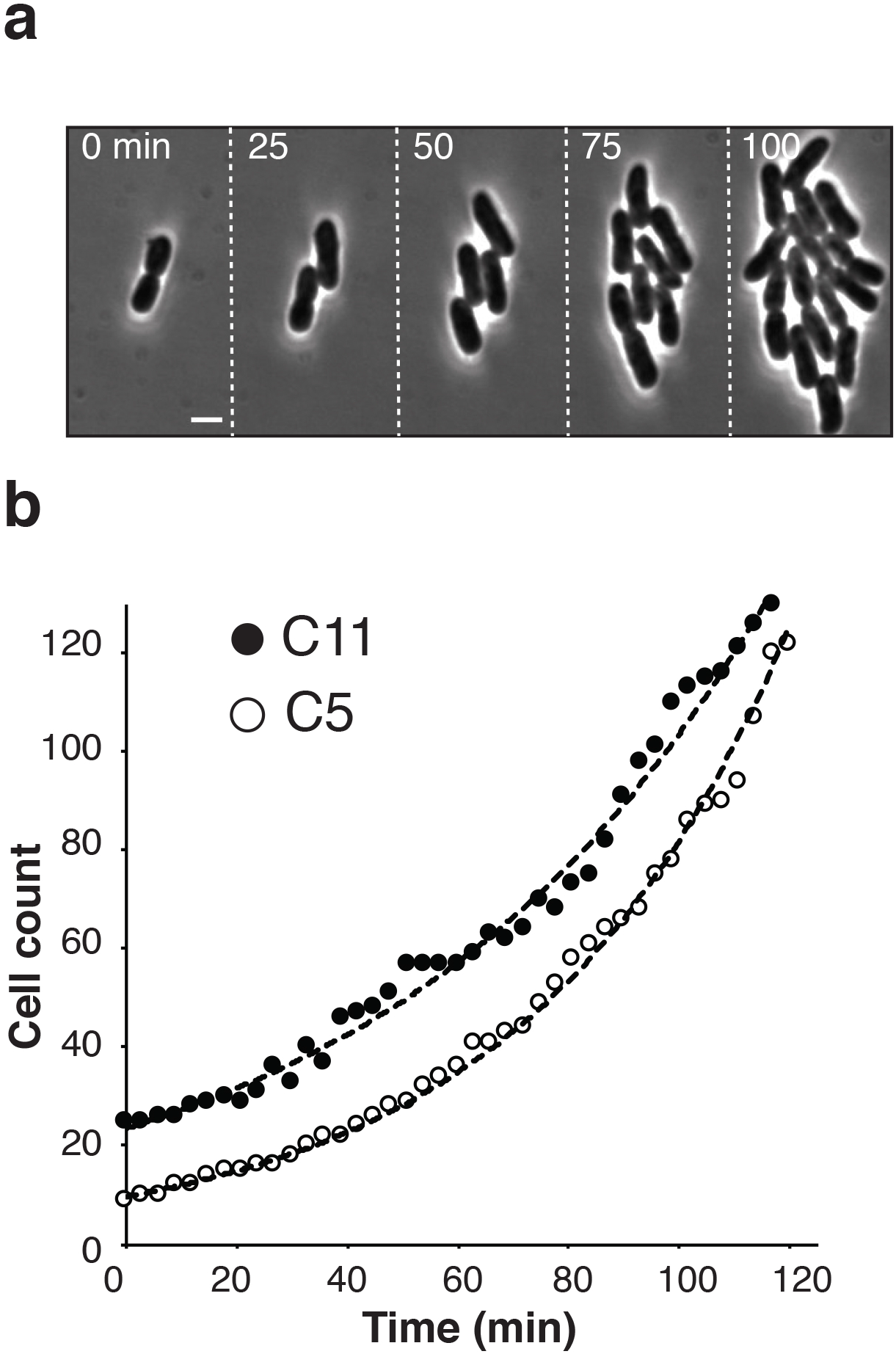
*Klebsiella pneumoniae* grows on CCS. **(a)** Growth of *Klebsiella pneumoniae* on C5. Shown are snapshots separated by 30 min after growth initiation (left panel). See associated Supplemental movie S5 for the full time-lapse. **(b)** Growth of *Klebsiella pneumoniae* on C5 and C11. Number of cells as function of time and corresponding exponential fits are shown.

We next wondered if additional “Klebsiella-compatible” chitosans could be identified. As discussed above, most tested chitosan polymers are not compatible with *E. coli* K12 and it could be interesting to identify “species-specific” polymers for AST. To do this, we further screened the Chito-library and successfully identified one additional CCS type, C11 (DA of 35 %, Mw of 557 kDa and thickness of 101 nm) that also supported *Klebsiella* adhesion and growth without detectable effect on bacterial fitness (Figure 4B, Figure S4). Importantly, C11 did not support growth of *E. coli*. In total, the results suggest that CCS is adaptable to the study of multiple bacterial species and that depending on their chemical structure chitosan substrates can either be derived to support adhesion and growth of multiple bacterial species or more specifically, to grow a given bacterial species or even perhaps strain.

## Discussion

In this work, we report a new glass functionalization procedure that supports bacterial adhesion and growth without any detectable physiological stress, contrarily to currently used polycationic polymers such as Poly-lysine. This technique allows studies of bacteria at the single cell level in simple microfluidic devices without the need of complex lithography or alternative physical immobilization techniques. Because the chemistry of the chitosan polymers and glass functionalization procedures are well established, the method is robust and highly reproducible. Moreover, CCS are long-lived and their integrity is not altered after storage of up to 6 months. Thus, CCS are highly versatile and provide a viable alternative to other and often more technically challenging microfluidic single cell approaches.

Although we characterized one CCS type in details and showed its potential for studies of *E. coli* and *K. pneumoniae* for basic and clinical purposes, we also show here that CCS can be derived for the studies of multiple strains and species, in particular ESKAPE pathogens. We also performed preliminary tests of the ability of C5 in promoting growth of a wide range of Gram negative and Gram positive bacteria. In our hands, C5 supported growth of *Vibrio cholera*, *Myxococcus xanthus*, *Mycobacterium smegmatis*, *Pseudomonas aeruginosa* (C5 also supported *Pseudomonas* twitching motility) but cell adhesion for these species was arguably not optimal. Nevertheless, CCS could be optimized for these species by testing different C5 thicknesses or alternatively, by isolating other CCS-types as we performed for *Klebsiella pneumoniae*.

The variable effects of chitosan between species and even within species is not too surprising because the biological properties of chitosan can vary widely based on composition and formulation. For example, chitosan polymers of large size (> 550 kDa) and high degree of acetylation (>50%) are known to exert bacteriostatic effects on some bacteria *(11)*. In addition, adhesion likely depends on the surface properties of the bacteria. In *E. coli*, phenotypic and genotypic diversity is very wide *(20)* and thus it is possible that some isolates fail to adhere (albeit a minority) because they have different surface properties (for example if they carry particular LPS O-antigens). An interesting avenue for future developments will be to test whether composite CCS made from several chitosan polymers increase the array of species and strains that may be grown on a single type of slides.

The applications of CCS in the field of bacterial cell biology are evident as such technology supports studies of any cellular processes, cell division, but also perhaps for studies of more complex population structures such as micro-colonies, biofilms and communities. Using a collection of *E. coli* strains, we typically observe that the bacteria first proliferate in two dimensions which we have shown by RICM occurs due to tight adhesion. The bacteria eventually proliferate away from the surface when space becomes a limiting factor for proliferation (Figure 2B). However, we also observed that some *E. coli* strains colonize the entire surface in 2D and thus form a single layer biofilm (Figure S5, Movie S6). The formation of *E. coli* micro-colonies on a surface has been shown to depend both on adhesion strength and preferential adhesion of the polar regions (which we also observe here, Figure 2B, *(21)*). Thus, it is likely that expanded micro-colonies are obtained depending on adhesion strength. Screening conditions that support biofilm formation for a particular strain could be achieved by defining a compatible adhesion range, either by modulating the ionic strength of the medium and/or changing the chitosan thickness, molar mass and DA.

The search for rapid phenotypic assays to determine antibiotics susceptibility is now a global priority to save on the use of large spectrum antibiotics and limit the spread of multiple antibiotics resistance in hospitals *(22)*. In current clinical practice, AST is generally performed using semi-automated methods that measure growth in bulk cultures in liquid (i.e. VITEK, *(23)*) or solid media. These methods only yield MICs estimate and the more accurate methods (i.e. antibiotic gradients or E-tests *(24)*) are time-consuming and costly. Moreover, all of these phenotypic antibiotic susceptibility testing require from 18 to 24 hours to provide an estimate of antibiotic susceptibility. Single cell microscopy approaches are powerful alternatives because they measure MIC as well as MCBs directly, more precisely and sometimes in less than 30 min, for example in microchannel chips *(2, 25)*. This technology however suffers from important drawbacks linked to sophisticated manipulation and high species-specific use, making its generalization in clinical practice difficult. Also, this method precludes morphometry analysis because the bacteria are maintained in channels that directly constrain their shape. Direct morphometry analysis for rapid AST has shown promising results on bacteria embedded in agarose *(3)*. However, in this case, the antibiotics were added indirectly by diffusion through the agarose making it difficult to control the exact concentrations and potentially slowing their action. In this context, CCS could provide an interesting alternative as we have shown that it can be applied reliably for two major ESKAPE pathogens and it combines the advantages of both above approaches, allowing direct antibiotic injection and morphometric analyses. The CCS method is more sensitive and ~10-20 times faster than traditional plate assays (here 50 min). A machine-learning based computational approach appears promising to measure MICs in automated fashion. The method still needs testing at higher throughput, but the results establish a proof of principle that its application for MIC determination is feasible. In addition, one system that could exploit it directly, the so-called Accelerate Pheno System (APS, Accelerate Diagnostics) is currently being implemented in hospitals *(26, 27)*. Similar to CCS, APS relies on surface immobilization of bacteria on glass slides in microfluidic channels. However, in absence of more adapted coating, the APS uses poly-lysine and Indium Tin Oxyde (ITO) facilitated gel electro-filtration to immobilize bacteria (US Patent N°7341841B2). It is therefore likely that this procedure perturbs the bacterial cell envelope, affecting the proton-motive force *(6–8)* and thus downstream, the accurate measurement of MICs. Indeed, lowering the proton motive force can artificially result in increased antibiotic resistance for some classes of antibiotics and thus generate false results *(9)*. In the future, adapting CCS to support growth of most clinical pathogens could make this technology an interesting tool for AST.

## Material and Methods

### Materials

Chitosans with low degrees of acetylation (DA) and different molar masses (*M_w_*) were purchased from Mahtani Co.ltd: a medium molar mass CS (CS_156_: DA 1.0%; *M_w_* = 156.1 kg/mol, *Đ* = 1.78, batch 243)) and a high molar mass CS (CS_557_: DA 2.4%; *M_w_* = 557.2 kg/mol, *Đ* = 1.39, batch 114). They were reacetylated to DA ranging from 1 to 80% using a procedure previously described. *(21)* Acetic acid (AcOH), hydrogen peroxide (40% w/w), sulfuric acid (96% w/w), hydrochloric acid (HCl, 37%), propan-1, 2-diol and ammonium hydroxide were purchased from Sigma Aldrich. Sterile and non-pyrogenic water was purchased from Otec^®^. Silicon wafers (doped-P bore, orientation (100)) were purchased from Siltronix^®^ and glass coverslips (75 x 25 x 0.17 mm^3^ #1.5H D263 Schott glass) from Ibidi.

### Chitosan preparation

CS was subjected to filtrations in order to remove insolubles and impurities before any use. CS was first solubilized in an AcOH aqueous solution, followed by successive filtrations through cellulose membrane (Millipore^®^) with pore sizes ranging from 3 μm to 0.22 μm. CS was then precipitated with ammonium hydroxide and washed by centrifugation with deionized water until a neutral pH was obtained. The purified CS was finally lyophilized and stored at room temperature.

In order to investigate the effect of DA on the film properties, CS with various DA were prepared by chemical modification using acetic anhydride, for both CS of different molar masses *(14)*. CS was first dissolved in an AcOH aqueous solution (1% w/w) overnight. A mixture of acetic anhydride and 1,2-propanediol was then added dropwise in the CS solution for at least 12 h under mechanical stirring. The amount of acetic anhydride added was calculated according to the DA aimed. The final solution was finally washed and lyophilized in the same manner as after the filtration step. The DA of the different CS prepared was determined by ^1^H NMR (Bruker Advance III, 400 MHz). For CS_156_, the DA obtained are: 9.0 %, 14.5 %, 25.6 %, 35.3 %, 41.9 % and 52.2 %. DAs close to those obtained for CS_156_ were obtained for CS_557_: 8.0 %, 12.2 %, 21.5 %, 34.0 %, 45.3 % and 52.6 %.

### Film preparation

Silicon substrates and glass coverslips were cleaned from organic pollution using a piranha bath (H_2_SO_4_/H_2_O_2_, 7/3 v/v) heated at 150 °C for 15 min, and then rinsed with deionized water (resistivity of 18 MΩ.cm). They were then subject to ultra-sonication in deionized water for 15 min and dried under a flux of clean air. The substrates (glass or silicon) were then placed into a plasma cleaner (Harrick Plasma ^®^) for 15 min in order to generate the silanol groups at the surface for a better adsorption of CS polymer chains.

In the meantime, CS was solubilized overnight in a solution of deionized water (Otec^®^) with AcOH, under magnetic stirring and at room temperature. The amount of acid added was calculated in stoichiometry compared to amine groups available along the CS polymer chain. CS solutions with different concentrations ranging from 0.3% to 1% for CS_557_ and concentrations ranging from 0.5% to 2% for CS_156_ were investigated in this study.

The films were finally formed onto silicon substrate by spin-coating at 2000 rpm until the solvent evaporates completely (5 min). After spin-coating, films were stored 24 h at room temperature before being characterized (unless mentioned otherwise). Some of the films were finally rinsed in AcOH aqueous solution (pH 4) for 5 min so that only the adsorbed chains of chitosan remain on the sample; the samples were then immersed in a water bath and finally dried under a flux of clean air (thickness < 3 nm in all cases).

### Surface topography

*AFM*. The surface morphologies were carried out by atomic force microscopy (AFM) (CSI Nano-observer). AFM probes with spring rate close to 40 N/m were purchased from Bruker. The AFM images were processed using Gwyddion software.

### Thickness measurement

The film thickness was measured on the silicon wafers using spectroscopic ellipsometry. On glass coverslips, the measurements were carried out using profilometry on scratched films. The consistency of the results obtained by ellipsometry or using profilometer profiles independently of the substrate used for a given chitosan solution permitted to use the measurement by ellipsometry as reference.

The ellipsometer (SOPRA GES-5E) was set at an incident angle of 70°, very close to the silicon Brewster angle. At least three measurements were done on each film at different positions in order to verify the film homogeneity. Data were then processed using WINELLI (Sopra-SA) software. A Cauchy model was used to fit experimental data (cos *Δ*, tan *Ψ*), in the spectral range of 2.0-4.5 eV, depending on fits and regression qualities, to evaluate the thickness. The UV parameters *A* and *B* were respectively set to 1.53 and 0.002.

A mechanical profilometer (Veeco Instruments) equipped with a cantilever of 2.5 μm in diameter was used to measure film thickness on glass coverslips. For this purpose, the samples were previously scratched by tweezer to locally remove CS film. Data analysis was performed with VISION V4.10 software from Veeco Instruments.

### Wetting measurements

Contact angles were measured using a tensiometer (Easydrop, Kruss) kit out with a camera connected to a computer equipped with a drop shape analysis software. To put down the liquids drop on the surface, a Hamilton syringe of 1 mL and a needle of 0.5 mm of diameter were used. “Static” measurements correspond to the angle determined 10 seconds after water drop deposition.

### Strains, Cell Cultures and Media preparation

Strains used were either lab strains (K12) or clinical strains obtained from the Laboratoire de Biologie - Centre Hospitalier Martigues.

*E. coli* and *K. Pneumoniae* cells were grown in ion-adjusted Luria-Bertani (LB) medium until exponential phase (OD = 0.5 ± 0.1) and diluted in LB to an OD around 0.01. The LB media was prepared using 10 g/L bacto-casitone (BD, 225930), 5 g/L NaCl (Biosciences, RC-093), 5 g/L bacto Yeast extract (BD, 212750) and osmosed water supplemented with 0.46 μg/L MgCl_2_, 3.21 μg/L CaCl_2_, 5.02 μg/L ZnCl_2_ and 6.15μg/L KCl. The cell suspension was then directly added to microfluidic channels. Loaded microfluidic chambers were centrifuged 3 min at 1000 rcf (Eppendorf centrifuge 5430R) to maximize cell adhesion.

### Preparation of chitosan slides and microfluidic chambers

Microfluidic channels were prepared from commercially-available six channel systems (sticky-Slide VI 0.4, IBIDI) that were directly applied to the surface of chitosan-coated slides. The dried chitosan was rehydrated by addition de-ionised MilliQ water for at least 5 min.

After centrifugation, the microfluidic channels were connected to a syringe and a pump (Aladdin syringe Pump WPI). Remaining non-adherent cells were thus removed trough a rinse step: 1.5 ml rinse with a 1.5ml/min flow followed by 1.5 ml with 5ml/min flow. The work flow was set at 3ml/h. Adhesion strength was assessed by increasing the flow in the channel. The shear stress was calculated by the following formula, given by IBIDI: τ = η· 176.1 ·Φ were τ is the shear stress (dyn/cm^2^), η the dynamical viscosity (dyn.s/cm^2^) and Φ the flow rate (ml/min). In absence of data about LB dynamical viscosity, we hypothesize that it is close to cell culture medium which is around 0.0072 dyn.s/cm2. On C5, adhered cells resisted shear forces above 12.3 dyn/cm2, indicating that they were firmly adhered.

### Dyes, Antibiotics treatment and MIC determination

Propidium iodide (PI) is used as a DNA stain that cannot cross the membrane of live cells, making it useful to differentiate healthy cells from dead cells. *E. coli* cells were immobilized to C5 chitosan on microfluidic chamber in presence or absence of antibiotics. Immediately before acquisition, the channel was rinsed with LB supplemented with 3mg/L Ertapenem (Sigma Aldrich) and 50μL/ml PI (Sigma Aldrich, P4170).

For MIC determination different channels were prepared simultaneously with the same cell suspension. Antibiotics Ertapenem, Ampicillin (Sigma Aldrich), Mecillinam (Sigma Aldrich) and Oxofloxacin (Sigma Aldrich) were prepared at different concentrations (one channel contained only LB as a control) and added to each channel just before image acquisition (every 3 min for standard acquisition). The MIC was defined for the lowest antibiotic concentration that induced cell death/stasis.

### Microscope acquisition and Image manipulation

Images were acquired with a Nikon phase contrast microscope (TE2000) equipped with a motorized stage, a Nikon perfect focus system and 100X objective lens. For technical convenience, experiments were performed at 25°C, a condition that supported both *E. coli* and *K. pneumoniae* growth. Standard Image analysis were performed under MicrobeJ a Fiji-Plugin developed for the analysis of bacteria *(28)*.

### Reflection Interference Contrast Microscopy (RICM)

RICM was performed with a Zeiss Observer inverted microscope (Carl Zeiss, Jena, Germany) equipped with a Zeiss Neofluar 63/1.25 antiflex objective, a crossed-polarizers cube, and a C7780 camera (Hamamatsu, Tokyo, Japan) with an adjustable field and aperture stops. The source was an X-cite 120Q lamp (Exfo, Mississauga, Canada) coupled to a narrow bandpass filter (λ=546nm+/-12 nm).

### Image segmentation

Image segmentation procedures were developed in Python. In order to provide a streamlined analysis procedure, we used the parameter-free threshold setting algorithm “iso_data” from the scikit-image python package *(29)* to extract the contours of the bacterial cells.

For each contour, we then perform a singular value decomposition from the numpy library *(30)* to retrieve aligned and centered contours for each bacteria. We use defect analysis (provided by the opencv library) to detect the septum and split the contours. If the defects attributed to the septum are distant of less than 0.5 μm and their center is less than 0.3 μm from the cell center, the contour is considered to be composed of two cells, and is therefore split.

From the detected and split contours, we then extract relevant morphometric data:

- the contour area,
- the contour length or perimeter,
- the longer of the min area rectangle,
- the width of the min area rectangle,
- the circularity defined as 4*πA/l*^2^. It is equal to 1 if, the contour is perfectly circular, lower than 1 otherwise,
- the inverse of the aspect ratio of the enclosing rectangle, (width/length), always lower than 1,
- the ratio of the minimal rectangle area to the cell area, (which should be close to one for a wild-type rod-shaped cell).

### Image annotation and training

In order to constitute a training-set to apply supervised machine learning, we developed a web-based dashboard based on plotly-dash toolset (http://plot.ly/dash). The annotation tool allows classifying the detected contours in 5 categories: normal, divided, abnormal, dead and invalid. We annotated 7 assays corresponding to 8300 contours.

### Outlier detection

From the annotated contours, those marked as “normal” were used to train a single class scalable vector machine classifier provided by the scikit-learn library *(31)* More precisely we fit a OneClassSVM object over 75% of the annotated data and use the remaining 25% over the above defined morphometric data. The trained classifier is then used on all the detected data to remove invalid contours from the count on each image.

### Sensitivity criterion

For each assay, the growth rate *G* is computed by performing a linear regression of the logarithm of the number of detected bacteria versus time.

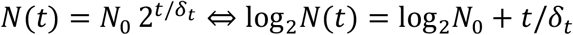

The reported error is the 95% confidence interval. We use the `scipy.stats.theilslopes` method *(32)* to perform the linear regression. A given growth assay is considered to survive if the growth rate of 0.2 *h*^-1^. This corresponds to a doubling time *δ* = *ln* (2)/*G* lower than 200 min. This cut off was chosen as it is longer than the microscopy acquistion span (Figure S3D).

## Acknowledgements

This work was funded by a CNRS Prematuration grant “Speedybiotics” and a SATT-Sud Est Maturation grant (Antiobio-R) to TM, OT and GS. We thank Leon Espinosa for help with the microscopy.

**Associated Supplemental Figure S1.**

**(a)** Procedure for chitosan purification and preparation (See Methods for details).

**(b)** Photographs of a silicon wafer before and after surface modification. In this example the wafer was covered with a chitosan solution of concentration 0.67%, DA 25% and *M_w_* = 180 kg/mol. The layer shows high homogeneity.

**Associated Supplemental Figure S2.**

**(a)** Morphological aberrations on Chitosan. Shown is *E. coli* K12 growth on C8 (Table 1). Note that the cells show abnormal twisted filaments that gradually detach over time. Scale bar = 2 μm.

**(b)** RICM of *E. coli* on torsion-inducing chitosan. On C8, RICM reveals that E. coli K12 cells are mostly attached by the cell poles, explaining their twisted shape due to the combined growth and local adhesion points. Scale bar = 4 μm.

**(c)** C5 shows no long-term toxicity. Shown are *E. coli* K12 cells that resume growth after being left for three days in exhausted medium and addition of fresh medium. Cell growth resumes normally. Scale bar = 8 μm.

**Associated Supplemental Figure S3.**

**(a)** Cell contour detection of a dividing bacterium after imaging by 100X Phase Contrast microscopy.

**(b)** Quantification of the extracted contour. The orange outline reveals the convex Hull of the detected contour. The red and gray dots are convexity defects used to detect the septum. Both red dots are automatically assigned to the septum. Further quantifications were based on the ratio of the area corresponding to the bacterial contour to the area of the minimal rectangle encompassing this contour (rectangular fill).

**(c)** Detection training of abnormal (and thus antibiotic sensitive) cells. Shown is an outlier detection obtained with the Scikit-learn one class scalable vector machine from the annotated dataset. For clarity, we only display the data corresponding to two dimensions, circularity and rectangular fill (see Methods for other parameters). Blue symbols: classified as normal; red symbols: classified as abnormal. Overall, there is a 10% rate of false negatives (normal contours labeled as abnormal) in both the training and test sets. This performance induces a noise is the growth curves and globally results in a 10% uncertainty in the computed growth rate. This uncertainty (which could be reduced with enhanced segmentation procedures) does not significantly affect the accuracy of MIC measurements.

**Associated Supplemental Figure S4:**

Growth of *Klebsiella pneumoniae* on C11. Shown are snapshots separated by 30 min after growth initiation (left panel). Scale bar = 2 μm.

**Associated Supplemental Figure S5: Single layer colonization of C5 by *E. coli* clinical strains.** Shown is the monolayer development of UTI698. Time points are separated by 54 min. Scale bar = 2 μm. See also associated movie S6

**Tables**

**Table 1. chitosan types and Adhesion and proliferation of *E. coli* K12**

**Table 2. MIC determination in clinical strains**

**Supplemental movies:**

Movie S1: Growth of *E. coli* K12 on C5

Movie S2: RICM of *E. coli* K12 on C5

Movie S3: *E. coli* K12 in the presence of Ampicillin

Movie S4: E. coli UTI 227 in the presence of Ertapenem and PI

Movie S5: K. Pneumoniae growth on C5

Movie S6: E. coli monolayers on C5

## References

1. D. I. Cattoni, J.-B. Fiche, A. Valeri, T. Mignot, M. Nöllmann, Super-resolution imaging of bacteria in a microfluidics device, PLoS ONE 8, e76268 (2013).

2. Ö. Baltekin, A. Boucharin, E. Tano, D. I. Andersson, J. Elf, Antibiotic susceptibility testing in less than 30 min using direct single-cell imaging, Proc. Natl. Acad. Sci. U.S.A. 114, 9170–9175 (2017).

3. J. Choi, J. Yoo, M. Lee, E.-G. Kim, J. S. Lee, S. Lee, S. Joo, S. H. Song, E.-C. Kim, J. C. Lee, H. C. Kim, Y.-G. Jung, S. Kwon, A rapid antimicrobial susceptibility test based on single-cell morphological analysis, Sci Transl Med 6, 267ra174 (2014).

4. A. Ducret, E. Maisonneuve, P. Notareschi, A. Grossi, T. Mignot, S. Dukan, A microscope automated fluidic system to study bacterial processes in real time, PLoS ONE 4, e7282 (2009).

5. Y. Matsumoto, S. Sakakihara, A. Grushnikov, K. Kikuchi, H. Noji, A. Yamaguchi, R. Iino, Y. Yagi, K. Nishino, A Microfluidic Channel Method for Rapid Drug-Susceptibility Testing of Pseudomonas aeruginosa, PLoS ONE 11, e0148797 (2016).

6. H. Strahl, L. W. Hamoen, Membrane potential is important for bacterial cell division, Proc. Natl. Acad. Sci. U.S.A. 107, 12281–12286 (2010).

7. T. Katsu, T. Tsuchiya, Y. Fujita, Dissipation of membrane potential of Escherichia coli cells induced by macromolecular polylysine, Biochem. Biophys. Res. Commun. 122, 401–406 (1984).

8. K. Colville, N. Tompkins, A. D. Rutenberg, M. H. Jericho, Effects of poly(L-lysine) substrates on attached Escherichia coli bacteria, Langmuir 26, 2639–2644 (2010).

9. B. Ezraty, A. Vergnes, M. Banzhaf, Y. Duverger, A. Huguenot, A. R. Brochado, S.-Y. Su, L. Espinosa, L. Loiseau, B. Py, A. Typas, F. Barras, Fe-S cluster biosynthesis controls uptake of aminoglycosides in a ROS-less death pathway, Science 340, 1583–1587 (2013).

10. S. Santajit, N. Indrawattana, Mechanisms of Antimicrobial Resistance in ESKAPE Pathogens, Biomed Res Int 2016, 2475067 (2016).

11. L. J. R. Foster, S. Ho, J. Hook, M. Basuki, H. Marçal, Chitosan as a Biomaterial: Influence of Degree of Deacetylation on Its Physiochemical, Material and Biological Properties, PLoS ONE 10, e0135153 (2015).

12. J. Nunthanid, S. Puttipipatkhachorn, K. Yamamoto, G. E. Peck, Physical properties and molecular behavior of chitosan films, Drug Dev Ind Pharm 27, 143–157 (2001).

13. A. Domard, A perspective on 30 years research on chitin and chitosan, Carbohydrate Polymers 84, 696–703 (2011).

14. L. Vachoud, N. Zydowicz, A. Domard, Formation and characterisation of a physical chitin gel, Carbohydrate Research 302, 169–177 (1997).

15. K. Drescher, J. Dunkel, C. D. Nadell, S. van Teeffelen, I. Grnja, N. S. Wingreen, H. A. Stone, B. L. Bassler, Architectural transitions in Vibrio cholerae biofilms at single-cell resolution, Proc. Natl. Acad. Sci. U.S.A. 113, E2066–2072 (2016).

16. L. M. Faure, J.-B. Fiche, L. Espinosa, A. Ducret, V. Anantharaman, J. Luciano, S. Lhospice, S. T. Islam, J. Tréguier, M. Sotes, E. Kuru, M. S. Van Nieuwenhze, Y. V. Brun, O. Théodoly, L. Aravind, M. Nollmann, T. Mignot, The mechanism of force transmission at bacterial focal adhesion complexes, Nature 539, 530–535 (2016).

17. A. D. Michelson, Platelets (Gulf Professional Publishing, 2002).

18. Z. Yao, D. Kahne, R. Kishony, Distinct single-cell morphological dynamics under beta-lactam antibiotics, Mol Cell 48, 705–712 (2012).

19. D. A. C. Stapels, P. W. S. Hill, A. J. Westermann, R. A. Fisher, T. L. Thurston, A.-E. Saliba, I. Blommestein, J. Vogel, S. Helaine, Salmonella persisters undermine host immune defenses during antibiotic treatment, Science 362, 1156–1160 (2018).

20. D. A. Rasko, M. J. Rosovitz, G. S. A. Myers, E. F. Mongodin, W. F. Fricke, P. Gajer, J. Crabtree, M. Sebaihia, N. R. Thomson, R. Chaudhuri, I. R. Henderson, V. Sperandio, J. Ravel, The Pangenome Structure of Escherichia coli: Comparative Genomic Analysis of E. coli Commensal and Pathogenic Isolates, Journal of Bacteriology 190, 6881–6893 (2008).

21. M.-C. Duvernoy, T. Mora, M. Ardré, V. Croquette, D. Bensimon, C. Quilliet, J.-M. Ghigo, M. Balland, C. Beloin, S. Lecuyer, N. Desprat, Asymmetric adhesion of rod-shaped bacteria controls microcolony morphogenesis, Nat Commun 9, 1120 (2018).

22. K. Sciarretta, J.-A. Røttingen, A. Opalska, A. J. Van Hengel, J. Larsen, Economic Incentives for Antibacterial Drug Development: Literature Review and Considerations From the Transatlantic Task Force on Antimicrobial Resistance, Clin. Infect. Dis. 63, 1470–1474 (2016).

23. T. K. W. Ling, P. C. Tam, Z. K. Liu, A. F. B. Cheng, Evaluation of VITEK 2 Rapid Identification and Susceptibility Testing System against Gram-Negative Clinical Isolates, Journal of Clinical Microbiology 39, 2964–2966 (2001).

24. L. B. Reller, M. Weinstein, J. H. Jorgensen, M. J. Ferraro, Antimicrobial Susceptibility Testing: A Review of General Principles and Contemporary Practices, Clin Infect Dis 49, 1749–1755 (2009).

25. J. Dai, M. Hamon, S. Jambovane, Microfluidics for Antibiotic Susceptibility and Toxicity Testing, Bioengineering 3, 25 (2016).

26. J. D. Lutgring, C. Bittencourt, E. McElvania TeKippe, D. Cavuoti, R. Hollaway, E. M. Burd, Evaluation of the Accelerate Pheno System: Results from Two Academic Medical Centers, J. Clin. Microbiol. 56 (2018), doi:10.1128/JCM.01672-17.

27. P. Pancholi, K. C. Carroll, B. W. Buchan, R. C. Chan, N. Dhiman, B. Ford, P. A. Granato, A. T. Harrington, D. R. Hernandez, R. M. Humphries, M. R. Jindra, N. A. Ledeboer, S. A. Miller, A. B. Mochon, M. A. Morgan, R. Patel, P. C. Schreckenberger, P. D. Stamper, P. J. Simner, N. E. Tucci, C. Zimmerman, D. M. Wolk, Multicenter Evaluation of the Accelerate PhenoTest BC Kit for Rapid Identification and Phenotypic Antimicrobial Susceptibility Testing Using Morphokinetic Cellular Analysis, J. Clin. Microbiol. 56 (2018), doi:10.1128/JCM.01329-17.

28. A. Ducret, E. M. Quardokus, Y. V. Brun, MicrobeJ, a tool for high throughput bacterial cell detection and quantitative analysis, Nat Microbiol 1, 16077 (2016).

29. S. van der Walt, J. L. Schönberger, J. Nunez-Iglesias, F. Boulogne, J. D. Warner, N. Yager, E. Gouillart, T. Yu, scikit-image contributors, scikit-image: image processing in Python, PeerJ 2, e453 (2014).

30. S. van der Walt, S. C. Colbert, G. Varoquaux, The NumPy Array: A Structure for Efficient Numerical Computation, Computing in Science & Engineering 13, 22–30 (2011).

31. F. Pedregosa, G. Varoquaux, A. Gramfort, V. Michel, B. Thirion, O. Grisel, M. Blondel, P. Prettenhofer, R. Weiss, V. Dubourg, J. Vanderplas, A. Passos, D. Cournapeau, M. Brucher, M. Perrot, É. Duchesnay, Scikit-learn: Machine Learning in Python, J. Mach. Learn. Res. 12, 2825–2830 (2011).

32. P. K. Sen, Estimates of the Regression Coefficient Based on Kendall’s Tau, Journal of the American Statistical Association 63, 1379–1389 (1968).

